# Assemblage of Focal Species Recognizers - AFSR: A technique for decreasing false positive rates of acoustic automatic identification in a multiple species context

**DOI:** 10.1101/546812

**Authors:** Ivan Braga Campos, Todd J. Landers, Kate D. Lee, William George Lee, Megan R. Friesen, Anne C. Gaskett, Louis Ranjard

## Abstract

Passive acoustic monitoring (PAM) coupled with automated species identification is a promising tool for species monitoring and conservation worldwide. However, high false positive rates are still an important limitation and a crucial factor for acceptance of these techniques in wildlife surveys. Here we present the Assemblage of Focal Species Recognizers - AFSR, a novel approach for decreasing false positives and increasing models’ precision in multispecies contexts. AFSR focusses on decreasing false positives by excluding unreliable sound file segments that are prone to misidentification. We used MatlabHTK, a hidden Markov models interface for bioacoustics analyses, for illustrating AFSR technique by comparing two approaches, 1) a multispecies recognizer where all species are identified simultaneously, and 2) an assemblage of focal species recognizers (AFSR), where several recognizers that each prioritise a single focal species are then summarised into a single output, according to a set of rules designed to exclude unreliable segments. Both approaches (the multispecies recognizer and AFSR) used the same sound files training dataset, but different processing workflow. We applied these recognisers to PAM recordings from a remote island colony with five seabird species and compared their outputs with manual species identifications. False positive rates and precision improved for all the five species when using AFSR, achieving remarkable 0% false positives and 100% precision for three of five seabird species, and < 6% false positive rates, and >90% precision for the other two species. AFSR’ output was also used to generate daily calling activity patterns for each species. Instead of attempting to withdraw useful information from every fragment in a sound recording, AFSR prioritises more trustworthy information from sections with better quality data. AFSR can be applied to automated species identification from multispecies PAM recordings worldwide.

## Introduction

Recent technical advances in sound-recording technologies and analyses have considerably enlarged the potential application of bioacoustics in conservation studies. Acoustic automated identification has been applied to numerous taxa including insects [1,2], anurans [3,4], bats [5,6], canids [7,8], birds [9–12], marine mammals [13] and elephants [14]. These automated techniques facilitate analysis of thousands of hours of sound files generated by a passive acoustic monitoring (PAM) approach, which could not realistically be done manually by a researcher. The integration of PAM recording with automated identification is a considerable advance that can be applied in biodiversity assessments in diverse environmental conditions and ecosystems. It can be particularly useful when species are rare, nocturnal or cryptic, or when sites are remote and have only intermittent access, e.g. seabird breeding colonies on offshore islands, which are a major conservation focus internationally [15].

Although these methods show great potential for wildlife monitoring, limitations and uncertainties remain that discourage incorporation of these analyses into management programs. Several researchers have identified high false positive rates [6,7,12,16–18]. These are typically associated with recordings involving multiple species [19], different environmental sound background [16], recording quality and overlapping calls [20]. Attempts to improve detection rates can also lead to more false positive identifications [5,18]. Given that passive acoustic monitoring is often the only method used to monitor species and populations, a low false positive rate is crucial.

Automated identification studies commonly focus on increasing detection rates in order to maximise the number of identified target calls in a sound file. However, in PAM, the amount of sound being recorded can easily reach terabytes of data. Potamitis et al. [21], for example, rejected 90–95% of their recordings because they did not meet their target specifications in the signal pre-processing stage. It is probably impossible to extract useful information from every single sound segment when monitoring long term. Instead, we advocate focus should be given to extracting more precise and trustworthy information from the segments with the highest probabilities of providing a true indication of species presence. In other words, automated identification models should prioritise low false positive rates instead of high detection rates and accuracy. Otherwise, even with adequate similarity and accuracy, a high false positive rate renders automated identification useless in a PAM context. Techniques for decreasing false positives are especially important in circumstances known to generate higher false positive rates, such as studies involving multiple species, which is a common situation in natural environments.

Another important factor for the utility of automatic recognizers is “precision”, the probability that a positive indication of presence is actually correct, or a true positive. Low false positive rates can be achieved by decreasing detection rates, but decreasing detection rates can also result in decreased true positives. When the rates of true positives and false positives are similar, even if both are low, any indication of a species presence would have similar chances of being correct or incorrect, so any technique that decreases false positives must also be reliable in terms of precision.

Here we present the Assemblage of Focal Species Recognizers - AFSR, a technique for decreasing false positive rates of automated identification models for acoustic recordings made in a multiple species context. It uses MatlabHTK, a hidden Markov models interface for bioacoustics analyses [22]. AFSR prioritises the extraction of information from more trustworthy sections of the recordings that can provide better quality data. It is a novel approach for increasing the precision of acoustic recognizers, increasing the applicability of automated identification techniques for wildlife surveys.

## Materials and Methods

### Study site and species

Pokohinu/Burgess Island (35° 54’ S, 175° 07’ E) is part of the Mokohinau archipelago, located ca. 90km northeast of Auckland’s east coast, in Aotearoa/New Zealand (Fig 1). The island became pest free after the eradication of the kiore or pacific rat (*Rattus exulans*) in 1990, and has since seen an increase in bird fauna [23]. We focus on the development of an automatic identification model for the calls of five Procellariiformes seabird species: *Pelecanoides urinatrix* (Kūaka, Common diving petrel), *Puffinus gavia* (Pakahā, Fluttering shearwater), *Pterodroma gouldi* (Ōi, Grey-faced petrel), *Puffinus assimilis* (Little shearwater), and *Pelagodroma marina* (Takahikare-moana, White-faced storm petrel). These species are known to have breeding colonies at Burgess Island, although they do not co-occur at all locations on the island [23].

**Fig 1.**
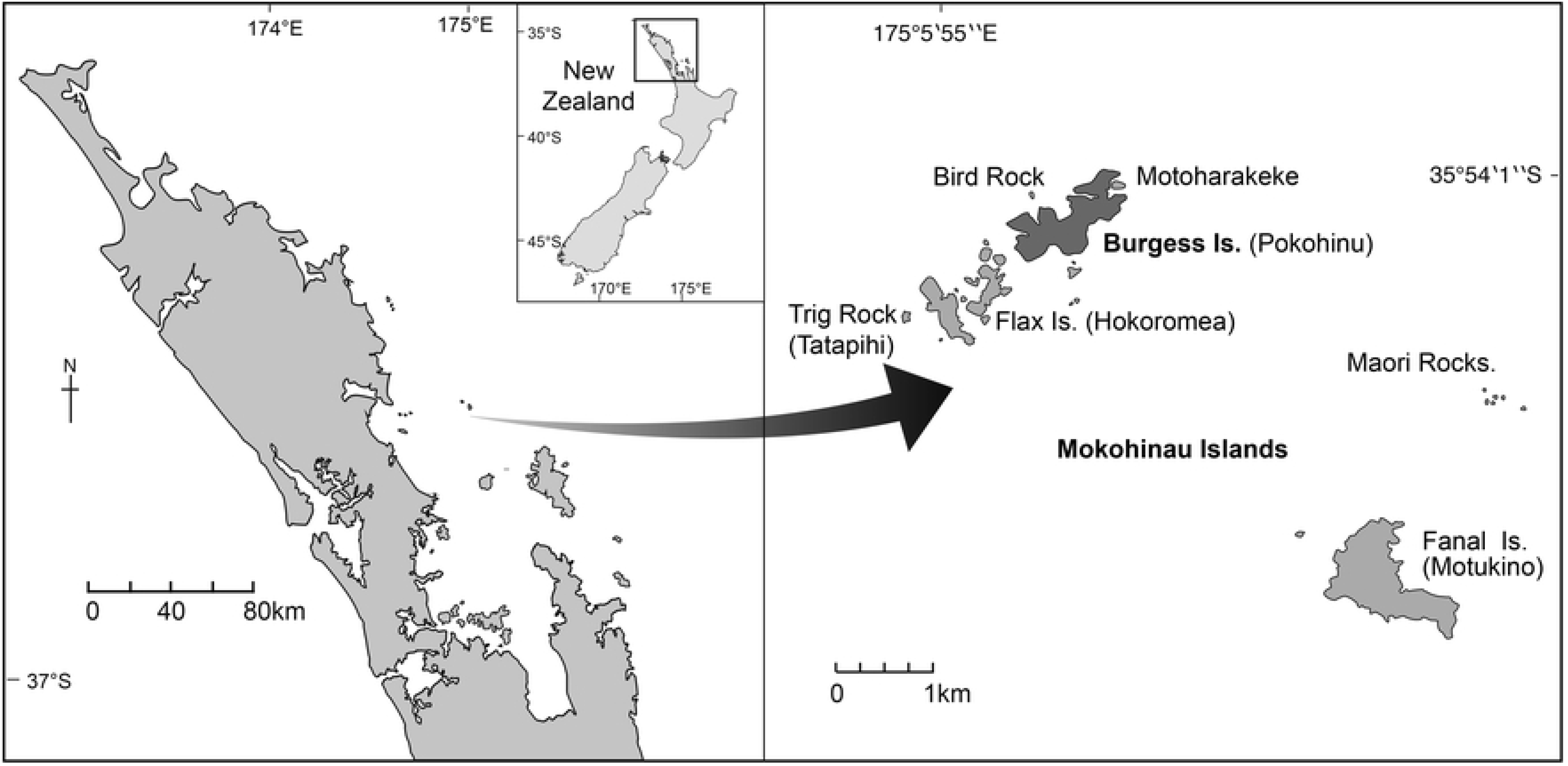
Pokohinu/Burgess Island, in the Mokohinau archipelago, Aotearoa/New Zealand.

### Acoustic Recordings

We used active recordings made by researchers in the field, and PAM recordings made without researchers’ presence using a pre-programmable acoustic sensor. The PAM recordings were made from 25^th^ - 30^th^ of September 2014, on Burgess Island (35.9057S, 175.1140E) using a sound recorder Song Meter SM2 (Wildlife Acoustics); at the sampling frequency of 44.1kHz; 32 bit resolution; from 6pm to 6am; totalling 60h of recordings. The active recordings were performed by seabird experts to obtain good examples of the birds’ typical calls, with confident species identification. Active recordings of Grey-faced petrels were made from April to late May 2015 at Bethells Beach, New Zealand, using a 722 Digital Audio Recorder (Sound Devices, LLC) with a Seinheiser highly directional microphone (model K6 ME 66; Wennebostel, Wedemark, Germany). Active recordings of the other four species were made on Burgess Island in September 2014 using a FR-2 Field Recorder (Foster Electric Co., Ltd.) with an Audio Technica shotgun microphone (model AT835b) housed in a Rycote wind-kit. All files were converted to 44.1kHz and 32 bit format.

### Building the training sound files data set

We built a data set of training sound files (total 179 MB) of good examples of the different species calls and their environmental background sound from sections of both the active and PAM recordings. From the active recordings, we selected and annotated calls that could be identified to species by seabird experts, and had good sound quality. We used these selected calls to create a preliminary species recogniser (S1 Supporting Information), that we then ran over the PAM recordings to fast track finding more examples of calls. All these calls extracted from the active and PAM recordings were combined into a single set of training sound files, which were then used for both our Multispecies Recogniser and our AFSR (assemblage of focal species recognizers). Even though the Multispecies Recogniser and AFSR were built from the same sound files, these files were associated with different sets of annotation text files in each approach and were processed through different workflows. An overall modelling workflow diagram is shown in the Fig 2.

**Fig 2.**
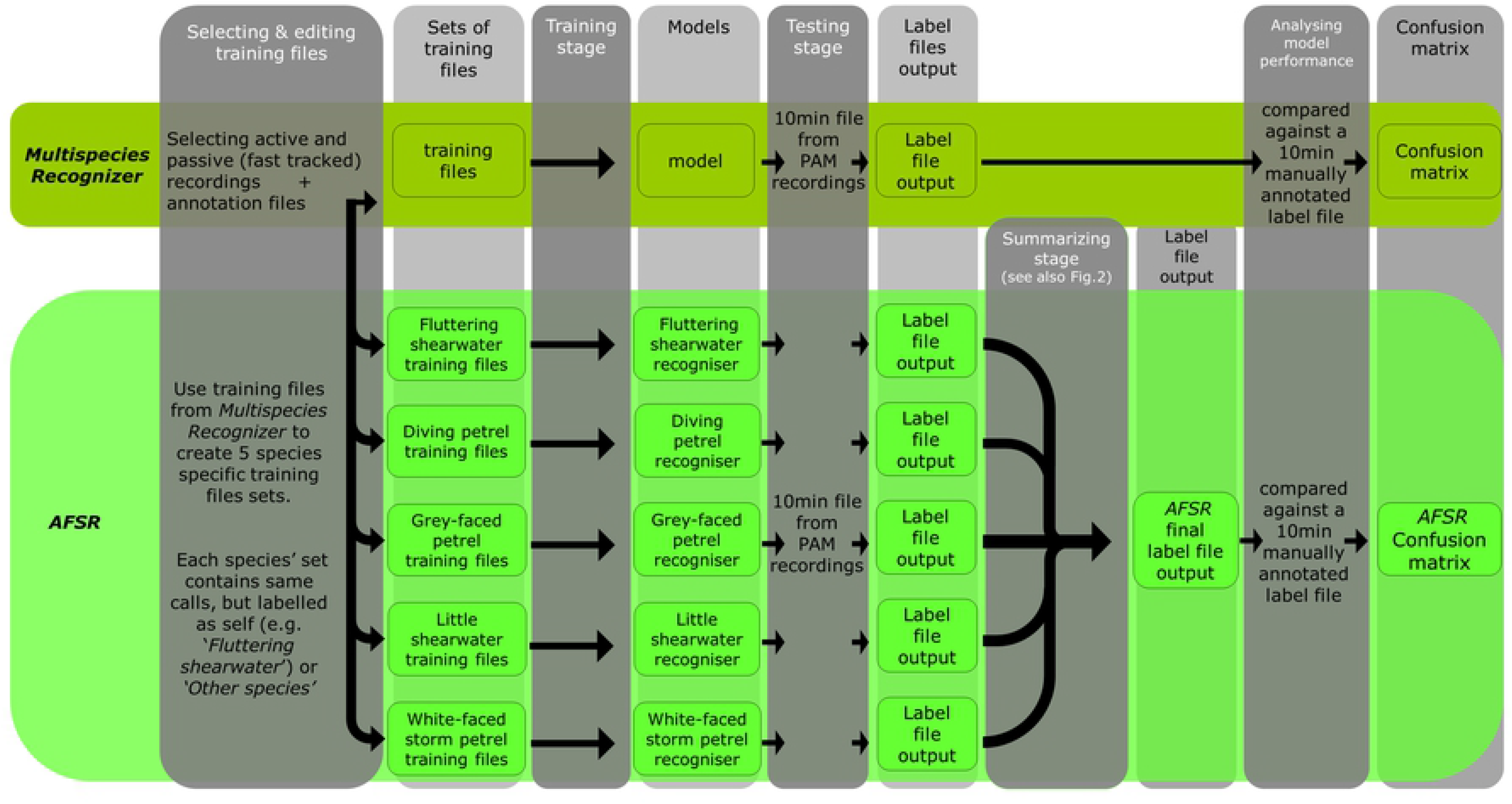
Modelling Workflow diagram. Each modelling approach is represented in horizontal lines. The workflow within each approach runs from left to right (columns) and the workflow from one modelling approach to the next runs from top to bottom.

#### 1) Multispecies Recognizer

Annotation files (“label” format) were manually created using the software AUDACITY^®^. These annotation files assigned a category name to each sound within the data set of training sound files described above. The categories used in the Multispecies Recogniser were: *Background; Diving Petrel; Grey-faced Petrel; Little Shearwater; Fluttering Shearwater;* and *White-faced Storm Petrel*. The ‘*Background*’ category was assigned to sound fragments where there were no existing petrel calls. This recognizer followed the same processing workflow as described by Ranjard et al. [22].

#### 2) Assemblage of Focal Species Recognizers - AFSR

Five independent species-specific recognisers were built using exactly the same data set of sound files data set previously described. In each case the sound files were associated with different annotation text files. For example, in the Little shearwater independent recogniser, all of this species’ calls were assigned in the annotation files as *‘Little Shearwater’*, while all the other four species’ calls were assigned as *‘Other Species’*. This framework was applied to all the five independent recognisers, one for each of our five seabird species. According to Potamitis et al. [21], a detection model needs to differentiate a target call from all non-target sounds, including other species’ calls and background sounds. In our system the category *‘Other Species’* was used in each one of the independent recognisers to help discriminate the target call from other species’ calls.

In this way, we obtained five species-specific outputs, which we then compared to detect and remove unreliable sections of the recordings prone to misidentification. We did this by creating a script named “AFSR_summarizing” that applies a set of rules to summarize the independent outputs into one final annotation text file. Whenever the five recognisers disagreed about the species identification of any segment of the sound recording, the section was then labelled as ‘*Unidentified*’. Only the recording segments that showed consistent species identifications by all five independent recognisers were considered a valid indicator of species presence. The data accessibility information containing the for “AFSR _summarizing” script and the link for the MatlabHTK package are presented on the S2 Supporting Information.

To this processing approach which consists in, from a single sound files data set, to create and run independent recognizers and then summarize their results into a single output following a specific set of rules, we named AFSR (Assemblage of Focal Species Recognizers). The set of rules used to summarize the five annotation text files (label format) into one is presented in the S3 Supporting Information. An illustration of the summarizing process is presented in the Fig 3.

**Fig 3.**
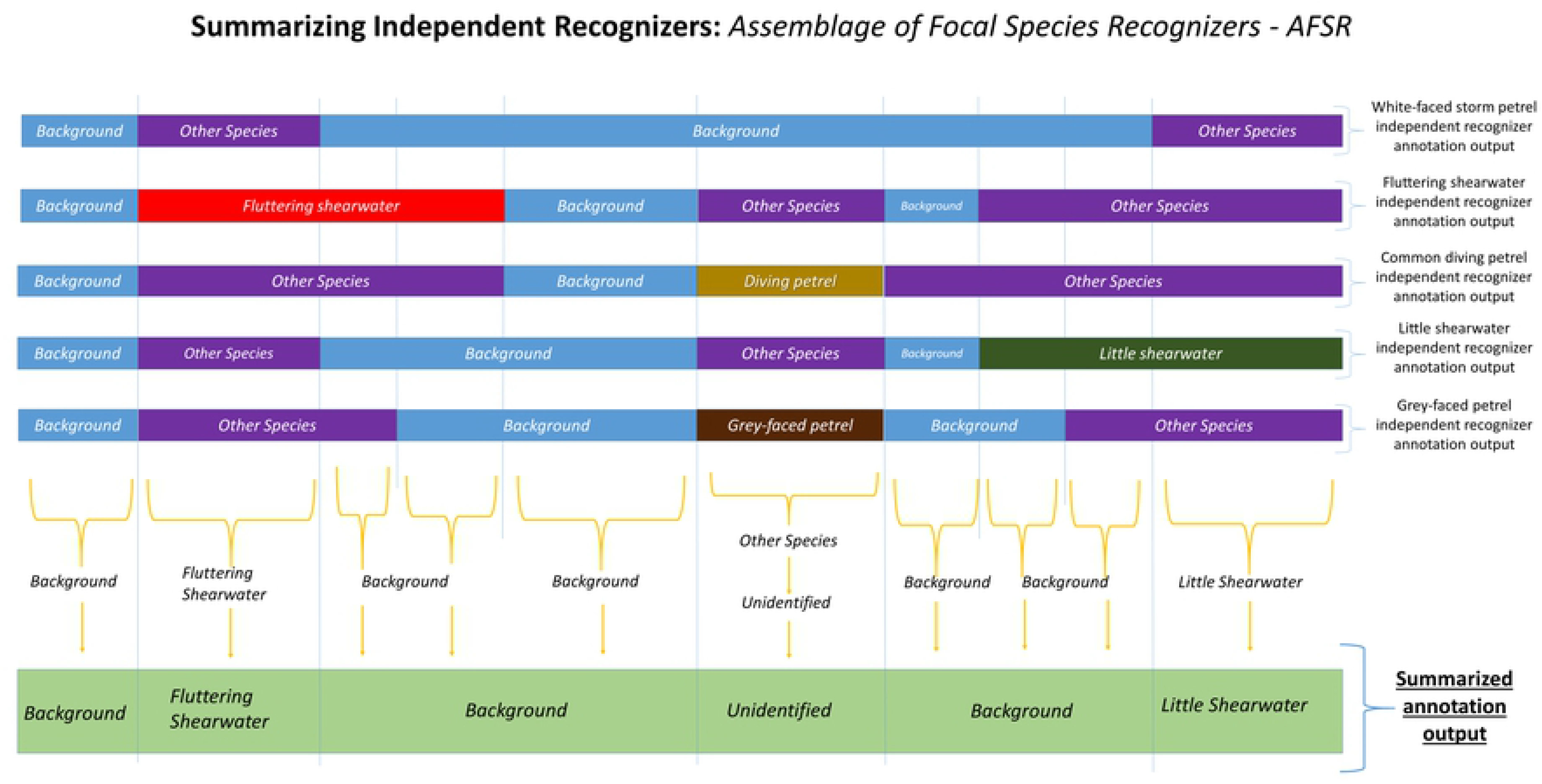
Illustration of the summarizing process. In this example of how AFSR converts category labels on a sound file fragment for five independent focal species recognisers into one summarized annotation based on specific rules, each of the upper horizontal lines represent one independent focal species recogniser’s annotation output and the lowest horizontal line represents the final summarized annotation output.

### MatlabHTK overview

The analyses were performed using the package MatlabHTK [22]. MatlabHTK processes the outputs of the Hidden Markov Toolkit (HTK; [24]), which was first designed for human speech analysis. In HTK, sound signal is typically represented as a sequence of parameter vectors, e.g. cepstrum coefficients, calculated for consecutive overlapping analysis windows. Hidden Markov Models (HMMs) statistically represent transitions between these windows. Therefore, HMMs are flexible and capable of dealing with variation of a given signal such as differing intensity, variations within species repertoire, diverse sound background, etc. MatlabHTK allows users to easily build and runs HTK acoustic models and to process the results.

Data analysis is typically performed in two stages; first, the HMM probabilities are estimated from training data (annotation text files + “.wav” sound files) and, second, recognition is performed on the data (sound files only). The recognition stage outputs annotation files that indicate which categories have been identified throughout the data sound files. A detailed description about the training and the recognition stages is presented by Ranjard et al. [22]. The signal processing and statistical analyses were performed through an Octave environment [25].

### Analysing the Recognizers

To assess the utility of the two modelling approaches, we created a 10 minute sound file containing calls of all the five species as well as background, which were extracted from Burgess Island PAM recordings. The seabird experts in our team listened to the 10 minute file to manually assign the correct species to every bird call in the sound file. The annotation of the 10 minute was made using 0.1 second time window (S1 Supporting Information), resulting in 6000 windows that were manually checked for species identification. Species identifications were verified by different team members, based on their extensive experience and knowledge of wild seabird calls. Additionally to the bird species, *‘Background’* was also used as a category assigned to background sound.

We then ran the Multispecies Recognizer and the AFSR over this 10 min file and compared their outputs with the manual identification made by researchers. A 0.4 second time window was selected for comparisons because this was the minimum length of any bird call in the training sound files (for a White-faced storm petrel call). To compare the two annotations, a similarity measure was defined: for each time window and for the two files, vectors indicating the percentage of the window’s duration assigned to each category were constructed. Then, the percentage similarity score “S” was defined as one minus the average Euclidean distance between vectors over all time windows [22]. Let w be the window size in seconds, c the number of categories, d^x^_1_ the duration in seconds of category x in the annotation file 1. Then, if the total number of windows is n, the similarity score between annotation file 1 and 2 is

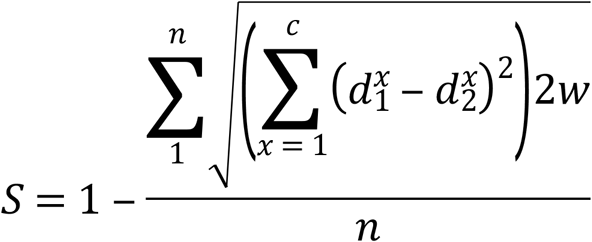

We generated a confusion matrix tables (“.csv” format) to compare the manually annotated file with the automatically annotated files (outputs) generated by the Multispecies Recognizer and AFSR. All the similarity scores and confusion matrices were calculated using the same 0.4 second time window. Precision scores for each target species was calculated from the confusion matrix values as follows:

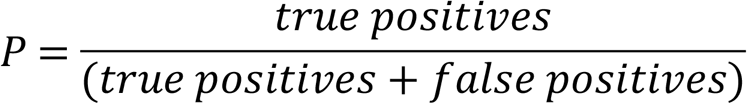

In a final step, to benchmark the biological relevance of the AFSR, we ran it over the five nights of PAM recordings and used the output to generate activity pattern data showing the time periods in which our five target species were actively calling at their shared colony on Burgess Island.

## Results

### 1) Multispecies Recognizer

This recognizer achieved an overall similarity of 82% when compared with the 10 minute manually annotated file. The false positive rates for four petrel species were lower than 10% (Little shearwater 5%; White-faced storm petrel 2%, Fluttering shearwater 1%, Common diving petrel 9%). However, the Grey-faced petrel had a higher false positive rate (10%). The precision was higher than 90% for three species: Little shearwater (93%), Fluttering shearwater (99%) and White-faced storm petrel (98%). However, the probabilities of false positives were > 10% for the Common diving petrel and Grey-faced petrel (precision was 88% for both species). The confusion matrix comparing the manual identification with the Multispecies Recognizer output is presented in the S4 Supporting Information.

### 2) Assemblage of Focal Species Recognizers - AFSR

The overall similarity score achieved by AFSR was lower than the Multispecies Recognizer (74% vs. 82%) but it considerably reduced the rates of false positives and increased precision scores for all five species. For White-Faced storm petrels, Fluttering shearwaters and Little shearwaters, the false positive rate achieved was 0% - all the indications of presence for these three species were correctly assigned. The false positive rates were also much lower for Common diving petrels (1%) and Grey-faced petrels (5%). For all species, precision values were >90%, with a remarkable 100% for Little shearwaters, Fluttering shearwaters, and White-faced storm petrels. Table 1 presents the total false positive rates and precision achieved by the AFSR and the Multispecies Recognizer. Table 2 presents the confusion matrix comparing the manual identification with the AFSR output.

**Table 1.** Total false positive rate and Precision per species achieved by Multispecies Recognizer and AFRS

| Species | Multispecies Recognizer |  | AFSR |  |
| --- | --- | --- | --- | --- |
|  | Total false positive rate | Precision | Total false positive rate | Precision |
| Common diving petrel | <u>0.09</u> | *0.88* | <u>0.01</u> | *0.97* |
| Grey-faced petrel | <u>0.1</u> | *0.88* | <u>0.05</u> | *0.92* |
| Little shearwater | <u>0.05</u> | *0.93* | <u>0</u> | *1* |
| Fluttering shearwater | <u>0.01</u> | *0.99* | <u>0</u> | *1* |
| White-faced storm petrel | <u>0.02</u> | *0.98* | <u>0</u> | *1* |

Total false positive rates (underlined) and precision (*italic*) for each species were calculated from the values generated by confusion matrices for each model and are presented here in a scale from 0 to 1, being 1 equals to 100%.

**Table 2.** Confusion Matrix comparing manual species identification Versus AFSR summarized output for a 10 minute long sound file

| Manually Annotated | AFSR |  |  |  |  |  |  |  |
| --- | --- | --- | --- | --- | --- | --- | --- | --- |
|  | Unidentified | Background | Common diving petrel | Grey-faced petrel | Little shearwater | Fluttering shearwater | White-faced storm petrel | Total false positive rate |
| Background | *0.08* | *0.75* | 0.04 | 0.01 | 0.03 | 0.01 | 0.08 | <u>0.17</u> |
| Common diving petrel | *0.33* | *0.29* | <u>0.37</u> | 0 | 0 | 0.01 | 0 | <u>0.01</u> |
| Grey-faced petrel | *0.19* | *0.21* | 0.04 | <u>0.55</u> | 0 | 0.01 | 0 | <u>0.05</u> |
| Little shearwater | *0.26* | *0.37* | 0 | 0 | <u>0.37</u> | 0 | 0 | <u>0</u> |
| Fluttering shearwater | *0.22* | *0.22* | 0 | 0 | 0 | <u>0.56</u> | 0 | <u>0</u> |
| White-faced storm petrel | *0.25* | *0.2* | 0 | 0 | 0 | 0 | <u>0.55</u> | <u>0</u> |

The proportion of the time in which each category indicated at the manually annotated text file is assigned to each one of the categories at the AFSR's output text file is presented in a scale from 0 to 1 (being 1 equals to 100%) as follows: cells with *values in italic*: negative indications of presence; underlined values: true positive indication of presence, values with no special formatting: false positive indication of presence; values underlined and italic: total false positive rate for each one of the categories (sum of the cells with no special formatting in each line).

AFSR results provided comprehensive daily activity patterns for individual species and the colony. Grey-faced petrels are the first species to vocalize at the colony after dusk (from ~19:20 hrs), shortly followed by Common diving petrels, Fluttering shearwaters, and White-faced storm petrels (all vocalizing from around ~19:30 hrs; Fig 4). The White-faced storm petrels are the first to depart before dawn around 04:20 hrs, followed by Grey-faced petrels and Fluttering shearwaters (about 5:00 hrs), then Common diving petrels (around 5:10 hrs). The low acoustic activity of Little shearwaters prevented a specific daily activity pattern from being produced.

**Fig 4.**
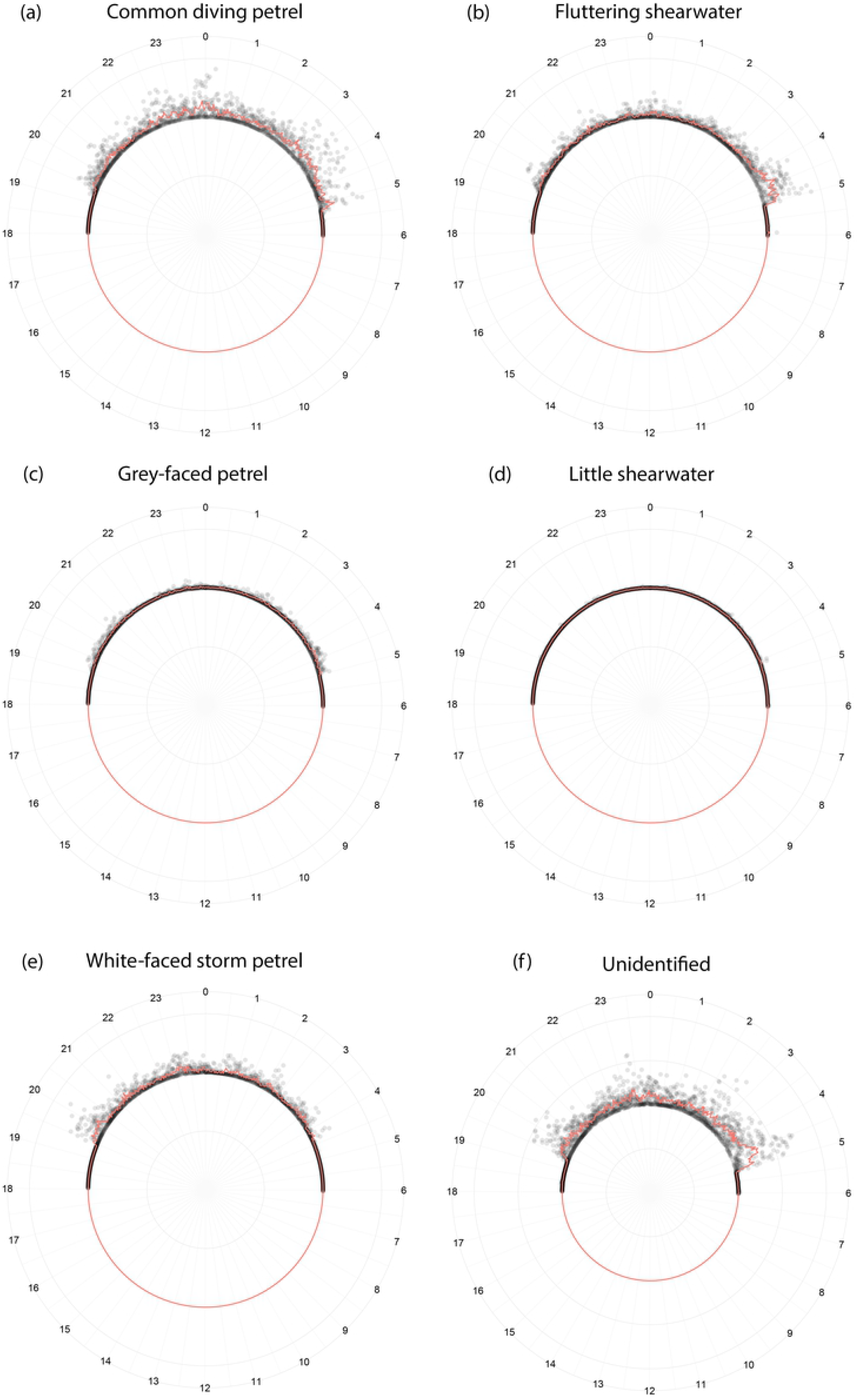
Daily pattern of acoustic activity identified by AFSR for the five seabird species. Common diving petrel [a]; Fluttering shearwater [b]; Grey-faced petrel [c]; Little shearwater [d]; White-faced storm petrel [e]; and the category Unidentified [f]; all identified from PAM recordings made for 5 consecutive days. Each grey circle is one indication of presence using 2 minute long analysis windows. The red line represents the average.

Fig 5 shows the average percentage of identification from 18:00 to 6:00 hrs for all categories combined (the five seabird species, as well as the *Unidentified* category). For the colony, the first peak of activity happens after the sunset when the birds arrive, from 19:20 to 20:30 hrs. The activity then declines but is persistent at some level through the night, with exception to the period between 23:00 and 00:00 hrs in which a moderate peak of activity occurs. The calling activity raises to the most intense level from 3:30 to 5:00 hrs, which occurs during the birds’ departure. After 5:00 hrs the activity reduces, and remains low until more birds arrive following the sunset. The overall acoustic activity pattern presented at Fig 5 is similar to the daily patter for calls assign to the *Unidentified* category (Fig 4-F).

**Fig 5.**
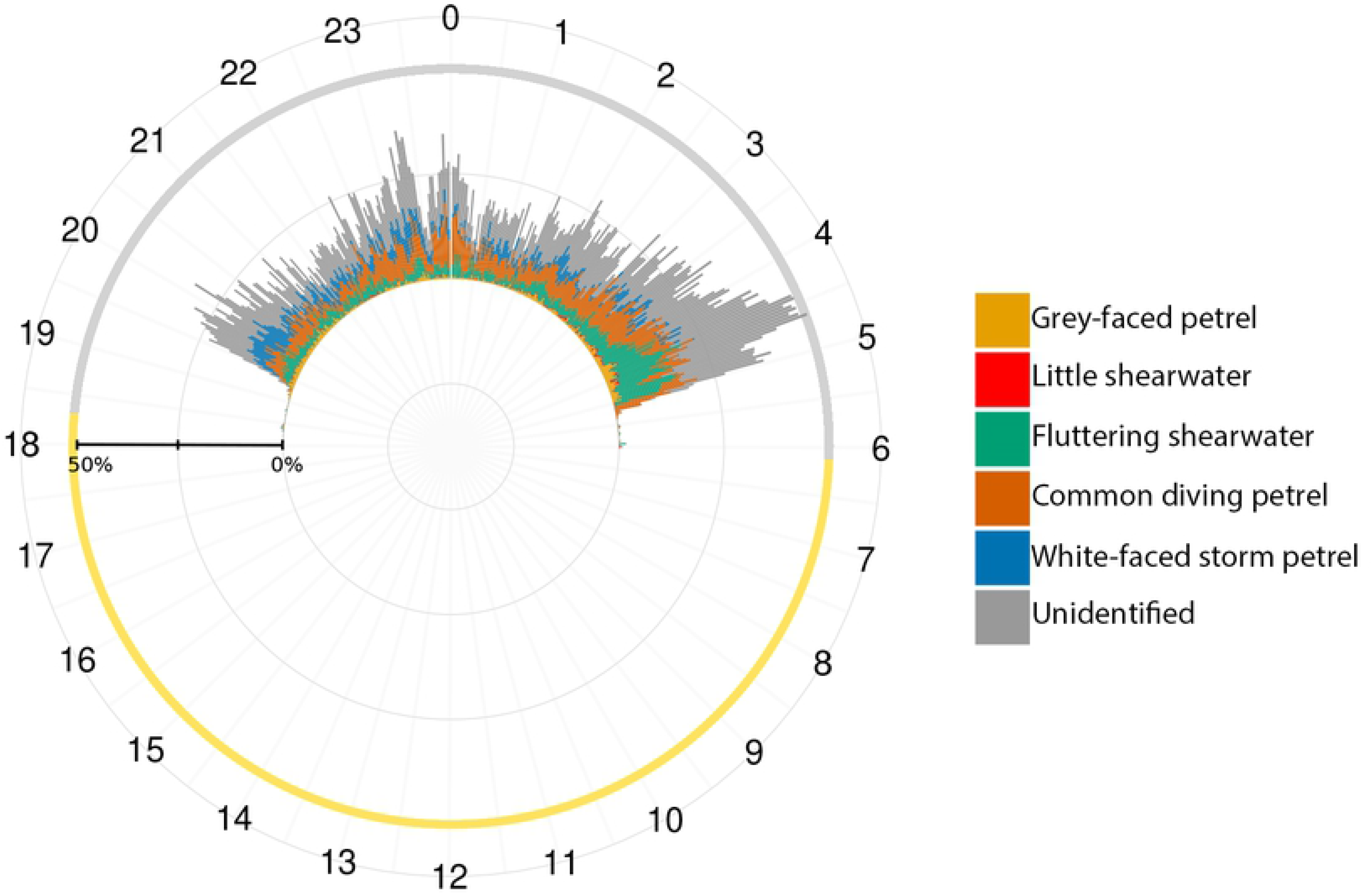
Colony mean daily acoustic activity for five seabird species, as well as sounds categorised as Unidentified by Assemblage of Focal Species Recognizers approach.

## Discussion

AFSR successfully decreased false positive rates and increased precision for all five target species in this comparative study of seabirds on an offshore island in New Zealand. Passive acoustic monitoring (PAM) recordings made in the natural settings can register complex soundscapes containing sound passages susceptible to generate misidentification. These sound passages can be a result of different individuals calling at the same time, call variations among individuals, different sound background, non-target species, among other reasons. When using the MatlabHTK interface [22], for every segment of sound in a testing file, only the category which best matches with the sound segment is indicated in the annotation output. The use of MatlabHTK combined with AFSR helps to significantly reduce false positives. The false positive and precision improvements achieved by AFSR are a result of the identification of the problematic sound fragments and their assignment to a new category (*“Unidentified”*), avoiding potentially incorrect indications of presence.

In order to prioritize reduced false positives and increased precision we accepted an overall decrease in similarity. As a consequence, we assumed we would have an increased level of false negatives. However, a relatively high false negative rate is not necessarily a problem for PAM. The three main sources of sound in a soundscape are biophony (sound produced by biological organisms); geophony (non-biological sound); and antrophony (human-induced noise) [26]. While geophony and antrophony are irrelevant for biodiversity monitoring, a portion of the biophony present in passive acoustic recordings is also not useful due to calls overlapping, signal masking, and other phenomena, which can produce poor quality sound segments. In general, when applying automatic identification to PAM recordings we should target sound segments with reliable and relevant information and leave poor quality remaining passages out of the analyses.

The use of AFSR is especially appropriate for multispecies monitoring at roosting and breeding colonies where the acoustic activity can be very intensive in some hours of the night, but absent at other times. This intense acoustic activity produces overlapping calls which makes their correct identification very difficult for conventional recognizer approaches and increases the chances of misidentification.

The biological relevancy of AFSR results is supported by the successful indication of presence of all the five species’ calls in our PAM recordings from Burgess Island. The activity patterns generated from the indications of presence confirm direct observations of these seabirds at natural colonies. The first and last peaks of activity (Fig 4) match the general pattern for petrels, with increased vocal activity just after sunset when birds arrive at the colony and before sunrise when they depart [27,28]. The first Grey-faced petrels’ calls recorded from around one hour after sunset, agrees with Ross and Brunton’s [29] data on arrival time at the colonies. The overall vocal activity pattern found for Common diving petrels is consistent with that reported by Ranjard et al. [22]. To our knowledge, there is no information about the daily activity pattern of the other studied species that we can use to compare to our automatically generated activity patterns. The low calling activity detected for Little shearwaters may be a consequence of a relatively smaller population size in comparison to the other four species at our fieldsite. Further studies are necessary to confirm this hypothesis.

It is important to highlight that the activity pattern of calls assigned to the *Unidentified* category (Fig 4 [f]) follows the overall acoustic activity pattern (Fig 5). This confirms our assumption that when more birds are calling, there is a higher chance of recording overlapping calls, and hence more misidentifications and false positives. These coincident patterns show our AFSR approach was effective in categorising calls that are problematic to identify and thus they were assigned to the *Unidentified* category. In this way, it reduces the false positives. Nevertheless, all the seabird species were detected by AFSR in this study, emphasising the success of this approach and its utility for multispecies monitoring.

Numerous islands in New Zealand have become pest free environments as a consequence of the elimination of invasive mammals which reduce or eliminate native bird’s populations [30]. Following pest eradication, many seabird re-establish breeding colonies on islands [31]. PAM associated with the *AFSR* model used in this paper can be an important tool for detecting and monitoring the process of the re-establishment of seabird breeding colonies in New Zealand islands and South Pacific region. However, the methods described here are likely to have broader applicability to any situation where multiple species with acoustic activity are being monitored to determine occupancy.

The use of AFSR also allows the addition of more species into a multiple species analysis. The same set of rules used for summarizing the annotations from the five independent models can be extended depending on the number of focal taxa. It makes AFSR suitable for being applied to multiple species from different animal communities and diverse ecosystems monitored by a passive acoustic monitoring approach around the world.

## Acknowledgments

IBC thanks CNPq-Brazil for financial support **through the** Science Without Borders scholarship program.

## Supporting Information

**S1 Supporting Information. Preliminary recognizers and overall modelling approach**.

**S2 Supporting Information. Data accessibility**.

**S3 Supporting Information. Summarizing rules**.

**S4 Supporting Information. Confusion matrix comparing manual species identification Versus the Multispecies Recognizer output for a 10 minute long sound file**.

